# Microbiome data enhances predictive models of lung function in people with cystic fibrosis

**DOI:** 10.1101/656066

**Authors:** Conan Y. Zhao, Yiqi Hao, Yifei Wang, John J. Varga, Arlene A. Stecenko, Joanna B. Goldberg, Sam P. Brown

## Abstract

**Background:** Microbiome sequencing has brought increasing attention to the polymicrobial context of chronic infections. However, clinical microbiology continues to focus on canonical human pathogens, which may overlook informative, but non-pathogenic, biomarkers. We address this disconnect in lung infections in people with cystic fibrosis (CF).

**Methods:** We collected health information (lung function, age, BMI) and sputum samples from a cohort of 77 children and adults with CF. Samples were collected during a period of clinical stability and 16S rDNA sequenced for airway microbiome compositions. We use Elastic Net regularization to train linear models predicting lung function and extract the most informative features.

**Results:** Models trained on whole microbiome quantitation outperform models trained on pathogen quantitation alone, with or without the inclusion of patient metadata. Our most accurate models retain key pathogens as negative predictors (*Pseudomonas, Achromobacter*) along with established correlates of CF disease state (age, BMI, CF related diabetes). In addition, our models select non-pathogen taxa (*Fusobacterium, Rothia*) as positive predictors of lung health.

**Conclusions:** These results support a reconsideration of clinical microbiology pipelines to ensure the provision of informative data to guide clinical practice.

## Introduction

Bacterial infections often resolve rapidly given effective immune responses, independent of antibiotic treatment. However, in chronic (long-lasting) cases, infections fail to clear even with appropriate drug treatment. Chronic infections impose an elevated morbidity and mortality risk to the individual [1] and an increasing burden on global healthcare systems as at-risk populations grow [2]. Chronic infections typically arise due to deficits in host barrier defenses and/or immune function, and commonly feature changes in pathogen growth mode (e.g. biofilm formation [3]) and additional microbial species acquisition, forming complex multispecies communities [4].

Microbiome sequencing has increasingly underscored the polymicrobial context of chronic infection. However, clinical microbiology analysis continues to focus only on the ‘usual suspects’ of established human pathogens – a relatively short list of organisms with well-established patient health risks. This disconnect between diverse ‘infection microbiomes’ and limited clinical microbiology profiling may overlook clinically important risk markers. To address this, we focus on chronic lung infections in people with cystic fibrosis (CF).

Cystic fibrosis is an autosomal recessive disease characterized by decreased lung mucociliary clearance and mucus accumulation [5–7]. The resulting environment provides both nutrients for bacterial growth and protection from host immune responses [8–11], facilitating chronic microbial infections [12–15]. Accessible 16S rDNA microbiome profiling has shifted CF airway microbiology research away from a historically single-pathogen focus, as sequencing expectorated sputum has revealed diverse communities of tens to hundreds of taxa, including numerous non-pathogenic bacteria [13,16,17].

Numerous lung microbiome studies have linked community composition to disease progression and overall patient health [18–20]. Cross-sectional studies have shown severe disease is associated with pathogen dominance and loss of taxonomic diversity [18,19,21]. Longitudinal studies have associated decreasing microbiome diversity with declining lung function [22]. Additionally, abundance of non-pathogenic fermentative anaerobes *(Veillonella, Prevotella, Fusobacterium)* is associated with higher lung function [23,24]. While these associations are observed across multiple studies, their causal interpretation is the subject of some controversy. These results may reflect community ecological processes within the lung, where species interactions govern community structure and subsequent harm to the host [12,25,26]. Conversely, these patterns could result from oral anaerobe contamination during sample collection [27,28]. Under this contamination model, increasing pathogen load compared to a constant background of oral microbiome contamination generates a spurious link between oral microbes, microbiome diversity, and patient health [27]. While recent paired sputum-saliva sampling analysis indicates that oral sample contamination is not a substantial contributor to sputum microbiome profiles in people with established CF lung disease [29], these conflicting hypotheses highlight the uncertainty in the role specific taxa present in sputum.

In the current study, we side-step this causal inference problem and instead assess how informative expectorated sputum microbiome data (including potential oral contaminants) is of patient lung health using a machine-learning framework. We hypothesize that the addition of non-pathogen data improves the prediction of patient lung function, compared to established pathogen data alone. To address this hypothesis we train predictive models on both lung microbiome and electronic medical record data for a cohort of CF patients. We find that compared to the benchmark of pathogen data alone, model performance was consistently improved by the addition of non-pathogen taxa.

## Results

### Clinical and microbiome data summary

In total, we obtained sputum expectorates from 77 CF children and adults. Pulmonary function, measured by percent predicted forced expiratory volume in 1 second (ppFEV1), was stratified into four categories from Severe to Normal. A summary of patient information is presented in Table 1. As expected, increasing age correlated with worsening lung function (ANOVA; p < 0.01). Culture-based detection of *Pseudomonas aeruginosa* correlated with decreasing lung function (ANOVA; p<0.001), as did (log-scaled) bacterial load (p < 0.05).

**Table 1.** Summary of patient clinical data, stratified by lung function. Lung function classes are defined as follows: Normal (ppFEV1 > 80); Mild (60 < ppFEV1 < 80); Moderate (40 < ppFEV1 < 60); and Severe (ppFEV1 < 40). Quantitative metrics are reported using the median and ranges. *Median reported values. **Two patients did not have reported HbA1c values. Significant differences between lung function categories tested by ANOVA, p-values shown. BC: Burkholderia, AX: Achromobacter, STE: Stenotrophomonas

|  | Severe | Moderate | Mild | Normal | P |
| --- | --- | --- | --- | --- | --- |
| **N** | 14 | 25 | 15 | 23 |  |
| **ppFEV1\*** | 32.9 | 46.5 | 74.9 | 101.2 |  |
| **(RANGE)** | (19.7-39.2) | (40.8-59.6) | (61.6-79.8) | (80.4-119.5) |  |
| **Age\*** | 31.5 | 32 | 24 | 20 | 0.007 |
| **(RANGE)** | (21-61) | (10-63) | (17-51) | (9-66) |  |
| **Male** | 6 | 11 | 7 | 11 |  |
| **CFTR Genotype** |  |  |  |  |  |
| **Homo-dF508** | 5 | 12 | 7 | 13 |  |
| **Hetero-dF508** | 9 | 10 | 8 | 10 |  |
| **Other/other** | 0 | 0 | 3 | 0 |  |
| **BMI\*** | 19.43 | 20.73 | 22.23 | 21.51 | 0.094 |
| **(RANGE)** | (16.27-25.69) | (16.70-29.81) | (19.38-26.07) | (16.65-33.91) |  |
| **CF-related diabetes (%)** | 11 (78.6) | 14 (56.0) | 8 (53.3) | 6 (26.1) | 0.015 |
| **HbA1c\*** | 6.25 | 5.9** | 5.7 | 5.5 | 0.009 |
| **(Range)** | (5.3-11.9) | (4.9-8.4) | (5.0-7.6) | (5.1-7.1) |  |
| **Clinical Micro** |  |  |  |  |  |
| **PA (%)** | 10 (71.4) | 20 (80.0) | 10 (66.7) | 5 (21.7) | <1.1e-4 |
| **SA (%)** | 8 (57.1) | 12 (48.0) | 10 (66.7) | 16 (69.6) | 0.454 |
| **MRSA (%)** | 4 (28.6) | 6 (24.0) | 6 (40.0) | 4 (17.4) | 0.486 |
| **BC (%)** | 0 (0.0) | 1 (4.0) | 1 (6.7) | 0 (0.0) | 0.553 |
| **AX (%)** | 3 (21.4) | 1 (4.0) | 0 (0.0) | 2 (8.7) | 0.147 |
| **STE (%)** | 0 (0.0) | 1 (4.0) | 3 (20.0) | 2 (8.7) | 0.191 |
| **16S metadata** |  |  |  |  |  |
| **% pathogen** | 0.857 | 0.589 | 0.532 | 0.195 | 9.05e-5 |
| **% Non-pathogen** | 0.135 | 0.404 | 0.597 | 0.783 | 3.01e-5 |

The majority (>90%) of reads from our sequencing analysis mapped to one of 13 genera (Figure 1a), consisting of both recognized CF pathogens (red) and orally derived bacteria (black). *Pseudomonas* sequences accounted for 30.4% of all reads, and were detected in every patient sample. Other established CF pathogens (*Staphylococcus, Achromobacter, Haemophilus, and Burkholderia*) collectively represented 19.3%, while oral taxa account for over 45% (Figure 1). Total pathogenic and non-pathogenic taxa abundance were both found to vary significantly (p << 0.001) with lung function (Table 1).

**Figure 1.**
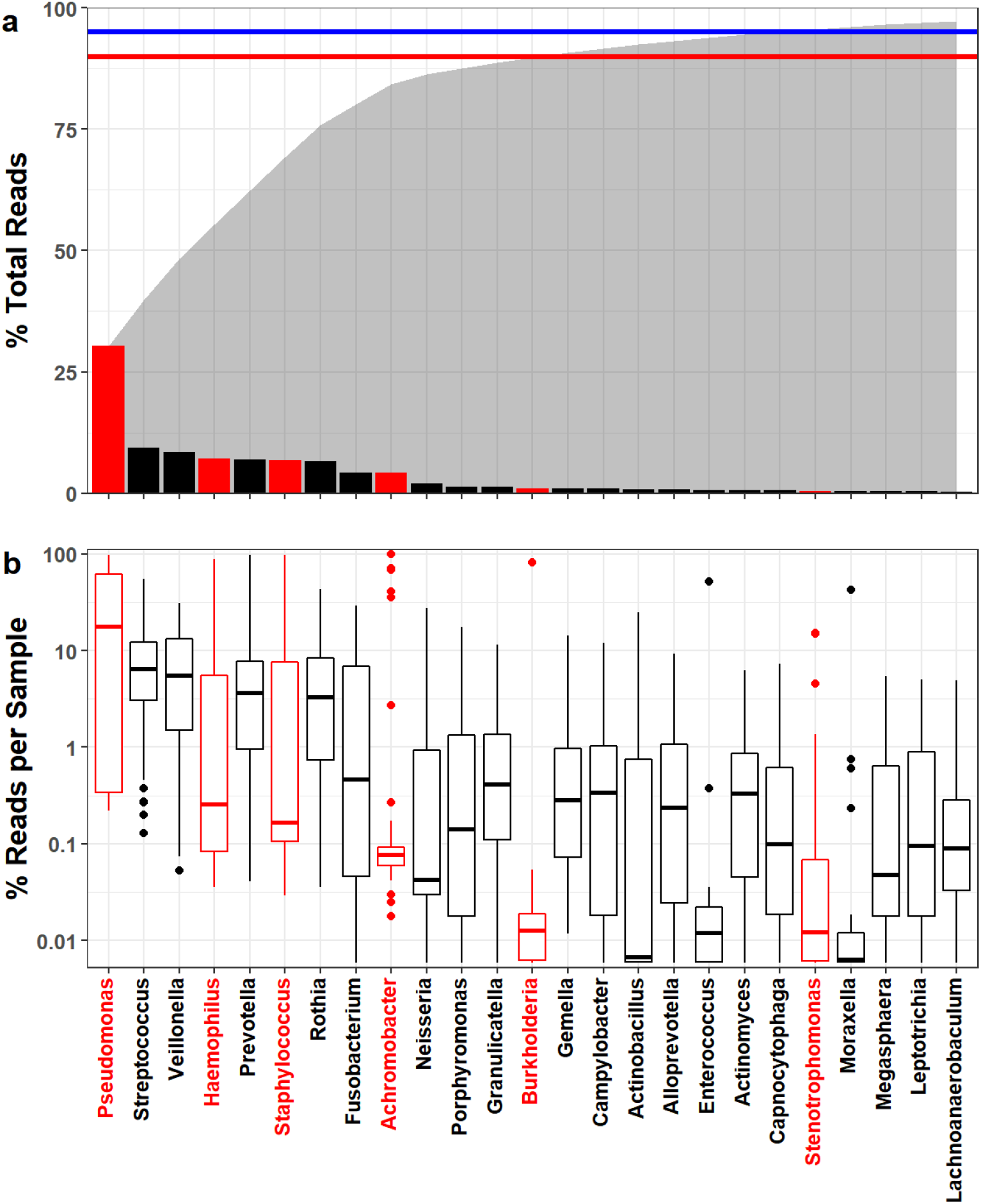
CF lung microbiomes are dominated by oral anaerobes and opportunistic pathogens. We analyzed CF sputum expectorate (N=77) using 16S sequencing and an in-house QIIME 2-based bioinformatics pipeline to resolve strain-level OTUs. Samples were rarefied to 17000 reads. We identified 217 OTUs across 59 genera and at least 81 species. Overall, we find that CF sputum samples are dominated by oral anaerobes and opportunistic pathogens. **a)** Sequences mapped to 14 genera comprised 90% (red line) of the total reads obtained. 95% (blue line) of all reads mapped to 21 genera. Total cumulative read fraction represented in shaded region. Pseudomonas was the most prevalent genus, followed by Streptococcus and Veillonella. **b)** Binning reads by sample shows variation in relative abundance. Pseudomonas comprises >10% of reads in the majority of our samples. While over 6% of the total reads mapped to Achromobacter, only 4 samples were comprised of >10% Achromobacter.

### Microbiome Composition Varies with Lung Function

We analyzed microbiome compositions across broad lung function categories to examine the relationship between sputum taxonomic profile and patient health. Figure 2a highlights the relative compositions of six canonical CF pathogens. As expected, *Pseudomonas* was more prevalent in lungs with reduced function, whereas in normal lungs *Haemophilus* and non-pathogen taxa (gray) were more prevalent. The non-pathogenic composition is consistently dominated by *Veillonella* and *Streptococcus* regardless of lung health or pathogen status (Figure 2b). Shannon diversity calculated with all taxa present is significantly greater for normal lung function (p<0.01, Figure S1a), in line with multiple other studies [30,31]. While Principle Coordinate Analysis (PCA) did not qualitatively separate compositions by lung function category, we found ppFEV1 was significantly associated with microbiome composition (Mantel test, r=0.195, p<0.001, Figure S1b).

**Figure 2.**
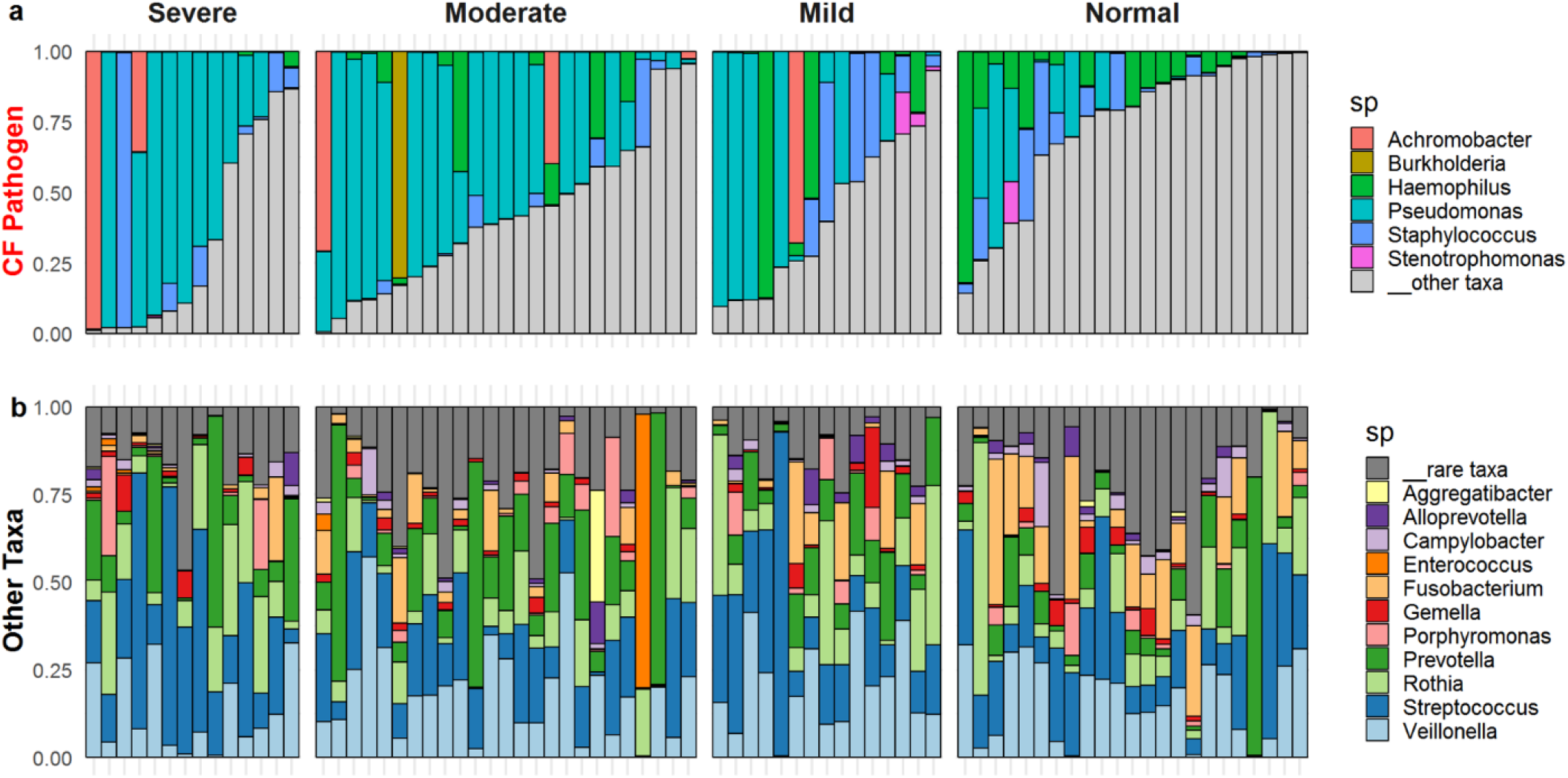
CF Lung microbiome composition varies with lung function and pathogen dominance. Relative abundances of **(a)** 6 canonical CF pathogens and **(b)** other taxa (the grey bar taxa in (a)). Microbiome compositions grouped by disease severity, classified by ppFEV1 score: normal (80+), mild (60-80), moderate (4060), and severe (<40).

### Integrating microbiome and patient meta-data

To examine multiple confounding variables such as patient age, BMI or CF-related diabetes (CFRD), we calculated spearman correlations across 14 microbiome, 11 patient metadata, and 6 clinical microbiology features (Figure 3). Hierarchical clustering reveals a complex autocorrelation structure, but with many expected consistencies. Overall, there are two main clusters of correlated variables. One correlated with ppFEV1, and included Shannon diversity index as well as 16S quantitation of *Fusobacterium, Haemophilus,* and *Neisseria.* The other anticorrelated with ppFEV1, and included ppFEV1 decline, pathogen abundance, CFRD and 16S quantitation of *Pseudomonas* and *Achromobacter.* Unsurprisingly, FEV1 and ppFEV1 cluster together and are inversely correlated with ppFEV1 decline rate (an average per year loss in ppFEV1 since birth). Additionally, 16S quantitation results for *Pseudomonas, Staphylococcus, Burkholderia,* and *Achromobacter* cluster with their respective culture-based clinical microbiology results. This does not hold for *Stenotrophomonas,* potentially due to its infrequent detection.

**Figure 3.**
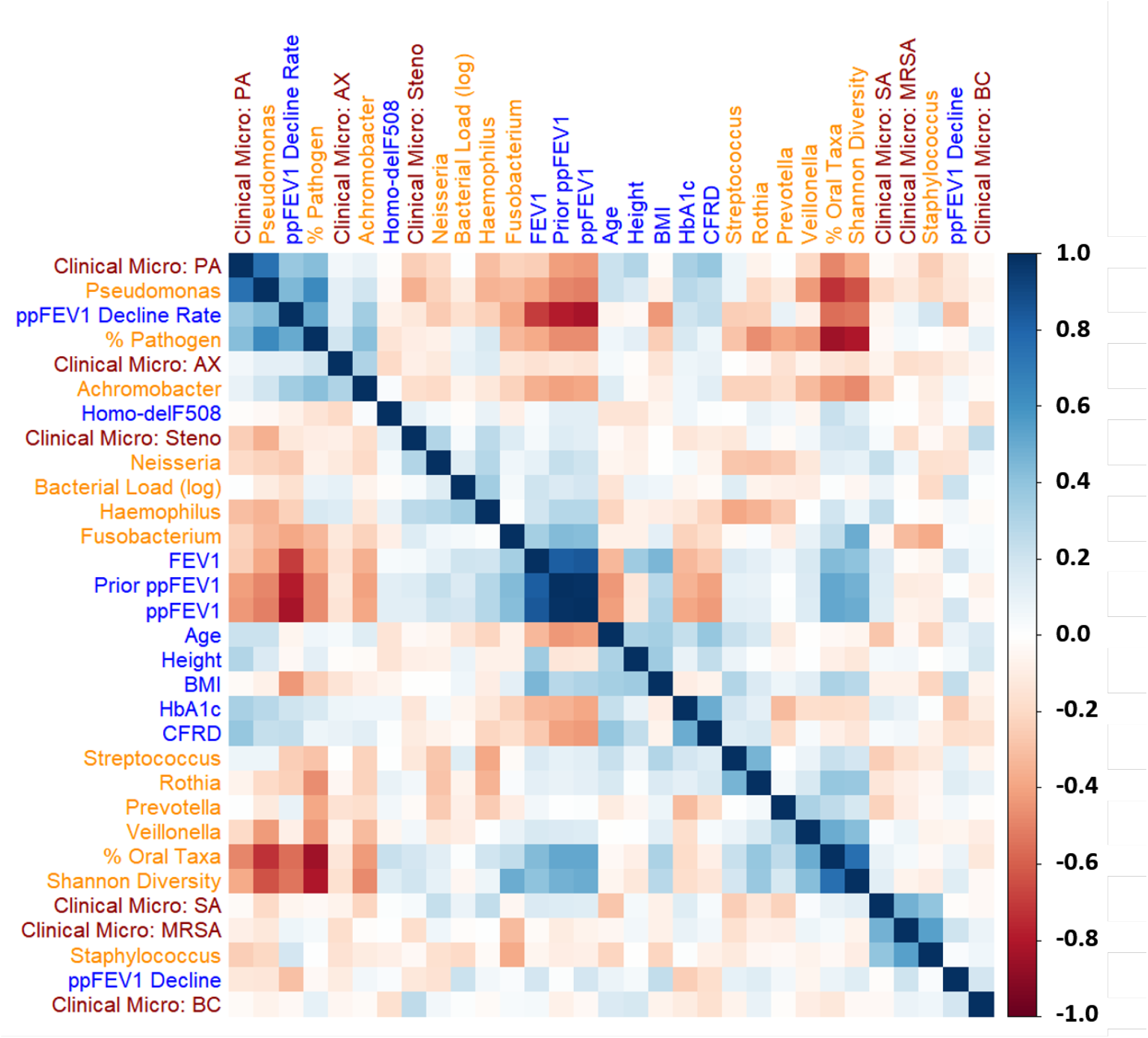
Lung function varies with patient meta-data. Spearman correlations (R::corrplot) across all patient metadata (blue), clinical micro results (maroon), and microbiome data (orange, clr-transformed) reveal a complex correlation structure. We used a centered-log transform on 16S data to mitigate compositional effects. Rows and columns were ordered by hierarchical clustering, which identified clusters of metadata and microbiome variables with similar correlation patterns.

### Dimensionality Reduction

The hairball correlation matrix in Figure 3 highlights the statistical challenges for identifying meaningful lung function predictors. Such challenges include high between-feature correlations and relatively few independent patient observations (N=77) compared to the initial number of available predictors (86 total, including 59 bacterial taxa). To mitigate this dimensionality problem, we first restrict our microbiome analysis to only the top 23 genera in our dataset. These top 23 encompass 97% of the total sequenced reads (Figure 1). We also calculate three additional summary statistics: % pathogen, % oral taxa, and Shannon diversity. As our clustering analysis shows reasonable agreement between clinical microbiology detection and rDNA sequencing, we exclude the binary detection results in favor of 16S quantitation. To address compositionality of 16S data, we incorporate total bacterial load (universal 16S primer qPCR) as a predictor. In addition, we use a centered logratio (clr) transform on our genus-level relative abundance data before standardizing to mean zero, unit variance inputs. We refer to this final combination of metadata and 16S data as our “All Features” dataset.

### Training Machine Learning Models

To assess if non-pathogenic taxa contain informative biomarkers, we split our samples into 53 training and 24 testing samples. ElasticNet was used to train predict lung function while performing feature selection (see methods, Figure 4). We expect that the addition of patient metadata (age, BMI etc) will improve our ability to predict lung function given the progressive nature of CF. Our null hypothesis, following the work of Jorth et al. and others[27,28] is that the taxa targeted by clinical microbiology provide adequate explanatory basis for lung function outcomes, and that the addition of non-pathogen 16S data will not improve model predictions.

**Figure 4.**
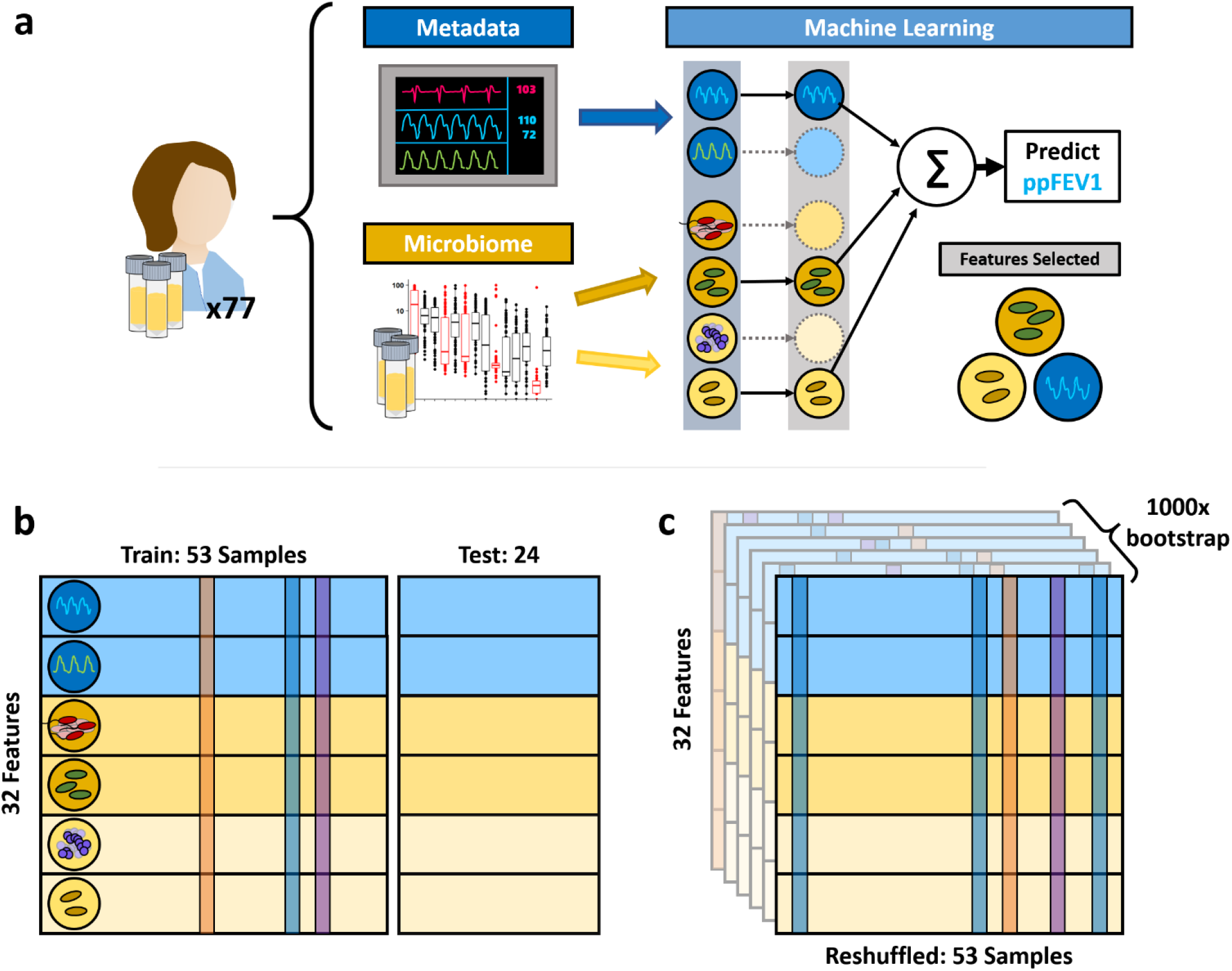
Machine Learning Overview. Machine learning models are trained on different input data tables using varying data resampling methods. (**a**) Features are categorized by information source (microbiome or patient metadata). The 16S data is further split into pathogens and other taxa in agreement with Figure 2. We use elastic net regularization to select informative features that predict ppFEV1. (**b**) We randomly selected 24 patient samples to withhold as a test set and train our models on the remaining 53 samples. To assess overfitting, we use leave-one-out cross validation on our training set. **(c)** We additionally implement 1000-fold bootstrap resampling to assess the robustness of our model fits.

We test this hypothesis by generating four additional feature subsets (CF Pathogens, All 16S Data, Metadata, and Metadata + Pathogens) and comparing the performance of models trained on each datasets. Initial-pass, non-bootstrapped model training results are shown in Figures S2 and S3.

### Model Generalizability

We assess overfitting using leave-one-out cross-validation and compare the prediction error across folds against the test set error. For model robustness, we use 1000-fold bootstrap resampling to fit both a baseline and ensemble of models. Robust features selected by the baseline model will also be selected by a large portion of the bootstrapped ensemble. We additionally standardize all features (mean=0, S.D.=1) to allow for crossfeature comparability. As an additional point of comparison, we generate a non-informative control dataset from the All Features set using within-feature shuffling, scrambling between-feature correlations while preserving the mean zero, unit variance within-feature structure. Figure 5 shows the results of our baseline (black points) and ensemble (boxplots) approaches. All models using patient metadata or microbiome data outperform the negative control.

**Figure 5.**
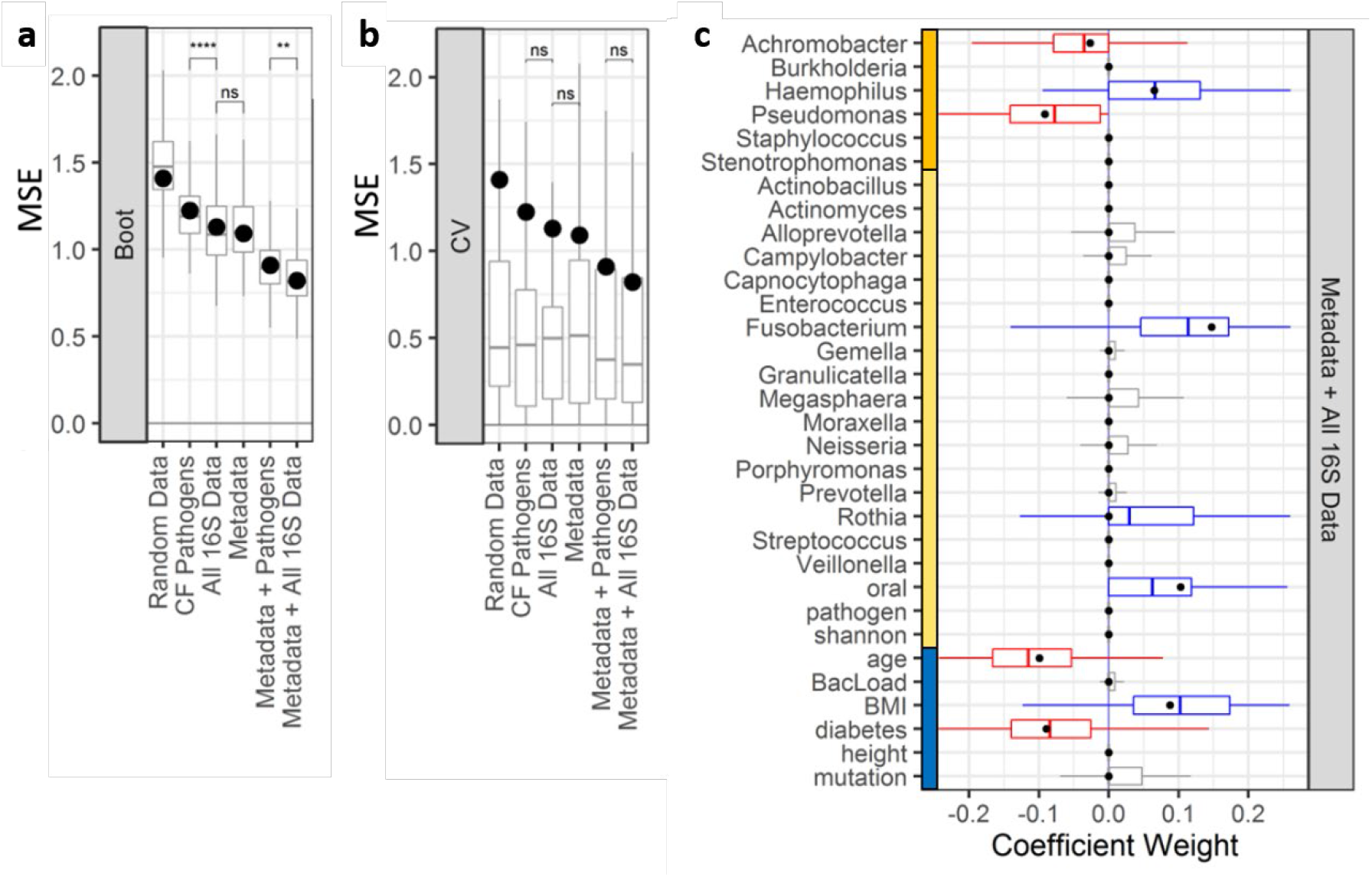
Bootstrapped ElasticNet-identified predictors of lung function. *ML models were trained using varying input datasets. **a)** 1000-fold bootstrapping and **b)** leave one out cross-validation (LOOCV) were used to generate prediction error (MSE) ranges across feature subsets. Models trained on all of the data show lower error compared to other feature subsets. Adding 16S pathogen quantitation decreases model error. Models trained on all 16S data outperform models using only 16S quantitation (p < 0.01, t test). Regardless of input features, models trained on the full sample set (black points) are greater than median LOOCV MSEs (boxplots). **c)** Coefficient ranges for train/test (black points) and bootstrapped models (boxplots) trained on standardized input datasets (blue: metadata, orange: 16S pathogens, yellow: 16S other taxa) show consistency between both machine learning strategies. Both cases select* Pseudomonas *and* Achromobacter *as negative predictors.*

### Addition of non-pathogen data improves model performance

To address the key question of relative model performance, we find that the addition of non-pathogen taxa significantly improves performance (significantly reduces bootstrapped MSE; Figure 5a), with or without the addition of patient meta-data. Models trained on all 16S quantitation significantly outperform models trained only on pathogen quantitation. Interestingly, while microbiome-only and metadata-only models achieve comparable performance, the combined model achieves greater model performance. Looking broadly across models, we find reasonable consistency in positive and negative predictor selection between our baseline and bootstrapped models (Figure S4).

We find multiple features selected across all training sets. *Pseudomonas, Achromobacter,* age, and diabetic status are consistently selected as negative predictors, while *Haemophilus, Fusobacterium, Rothia,* oral taxa abundance, and *BMI* are consistently positive predictors. All informative features selected in the independent models (Figure S4c) were also selected in the All Features model (Figure S4g). A small subset (< 50%) of the bootstrapped models also selected a handful of oral taxa, bacterial load, and CFTR mutation type as positive predictors of lung function (Figure 5c, gray boxplots). However, a majority of bootstrapped models and the train/test model did not select these as informative features.

As an additional check against overfitting, we obtain ranges of model errors (measured by mean squared error of predicted ppFEV1 values) using leave-one-out cross validation (Figure 5b). We do not find significant differences between cross-validated model errors across our training sets, suggesting that despite the difference in number of available predictors, our models are not overfitting.

## Discussion

People with CF face the challenge of managing long-term chronic infections. Current respiratory management practice is driven by clinical microbiology identification of specific pathogens in throat cultures or expectorated sputum samples, alongside measures of respiratory status (changes in symptoms, signs, and/or lung function). In the current study, we used 16S sequencing to assess sputum microbiome content more broadly, and ask whether the addition of non-pathogen taxa improves our ability to predict patient lung health, with or without the inclusion of patient health data. To address this question we applied machine learning tools to an integrated 77 patient lung microbiome and electronic medical record dataset. Our analysis revealed that the addition of non-pathogen data improves prediction of patient health, with the most accurate models selecting patient metadata, pathogen quantitation, and non-pathogen information. Our inclusive ‘all data’ models additionally point to a predictive role for specific non-pathogen taxa, in particular the oral anaerobe genera *Rothia* and *Fusobacterium.*

Despite the significant contribution of non-pathogen data, our results are still broadly consistent with what might be termed the ‘traditional’ view of CF microbiology. Established CF pathogens (*P. aeruginosa, S. aureus, H. influenzae, B. cenocepacia*) are the major drivers of patient outcomes, as evidenced by substantial improvement in predictive outcomes whenever we include pathogen data (Figure 5a), and the by comparison relatively weak contribution of the addition of non-pathogen taxa. Note that we specifically use quantitative 16S measures of pathogen composition to provide a level playing field in the comparison of pathogen and non-pathogen predictive contribution. Figure 3 highlights that quantitative 16S and qualitative (presence/absence) clinical microbiology data are in general agreement.

The traditional role of CF pathogens as the central predictors of patient outcomes has been challenged over the past decade by the advent of microbiome sequencing. Extensive surveys have documented an association between CF lung function and microbiome diversity, also evident in the current study (Figure 2). At face value, these results suggest a biological role for these non-pathogen taxa, potentially competing with [32] or facilitating [33] pathogen taxa and therefore indirectly shaping disease outcomes. Jorth et al. recently published a forceful rejection of this ‘active microbiome’ view, stressing the potential causal role of changing pathogen densities in shaping disease outcomes and viewing shifting diversity metrics as a simple statistical ‘relative composition’ artifact of shifting pathogen numbers against a roughly constant oral contamination background [27]. While our analyses provide some support for this view, in particular the constancy of the non-pathogen microbiome across patients (Figure 2b) and the lack of substantial predictive improvement on addition of non-pathogen data (Figure 5b), we also see lines of evidence against the contamination hypothesis. First, our use of center log transformations mitigates the risk of spurious associations due to compositionality (refs) and yet non-pathogen taxa are still consistently retained. Second, the contamination hypothesis predicts total bacterial burden to be an important predictor, and yet burden was not retained in our models. Third, our observation of a consistent retention of specific non-pathogen taxa across multiple models (with and without the addition of potentially confounding EMR features, including age and BMI) points to the potential for a distinct causal pathway that is orthologous to age or BMI. We note that the interpretation that oral bacteria are active players in the lung environment is further buttressed by a recent study on people with established CF disease (Lu et al. 2020) that used paired sputum and saliva samples to infer the presence of substantial populations of oral bacteria in the lung.

Our ‘all data’ models highlight *Rothia* and *Fusobacterium* as positive predictors of lung function across our 77 patients, in models that already take into account pathogen data. When we include features already known to correlate with lung health, such as age, BMI, and CFRD status, our models not only these features, but additionally retain *Rothia* and *Fusobacterium* as positive predictors. The retention of these specific taxa in both this full model and in partial models (Figure S4) suggests that these taxa provide potentially valuable predictive information on current patient health. Of course, this analysis does not allow inference to causal mechanism or even direction of causality. It is entirely possible that these taxa are simply bio-markers of dimensions of improved health that are largely independent of age, BMI, and other established positive predictors that are already accounted for in the model. It is also possible that these specific taxa play a more active causal role, for instance holding specific pathogens at bay via competitive interspecific mechanisms [34].

Interestingly, our All Features models also highlight *Haemophilus,* a canonical CF pathogen, as a positive predictor of lung function. *Haemophilus influenzae* infections are most common in younger CF patients [8,35], hence we would expect a positive association in a model that is not controlled for age (Figure S4c, S4d). However we see that the positive weighting on *Haemophilus* is retained in models that also account for age as a positive predictor of lung function. A second possibility is that the positive weighting of *Haemophilus* is due to pathogen-pathogen competition and the relatively less severe nature of *Haemophilus* infections in adults (i.e., *Haemophilus* is ‘best of a bad job’). Figure 2a illustrates that we only appreciably detect two and rarely three coexisting pathogens of the six we find across all patients. The relative scarcity of multi-pathogen communities implies that *Haemophilus* presence coincides with the absence of other more severe pathogens – and indeed we see a dominance of negative correlations among pathogens (Figure 3). In this context we cannot preclude a protective role of *Haemophilus* against more severe pathogens in older patients.

A caveat of this analysis is the dependency of machine learning performance and robustness on particular distributions of data, and the failure of linear algorithms such as LASSO and ElasticNet on microbiomelike data [36–38]. This is in part due to the compositionality constraint of microbiome data, which can be mitigated by using absolute quantitation [39]. However, training on absolute abundances introduces additional caveats, as order-of-magnitude differences in qPCR sample quantitation can in turn over-represent samples with higher bacterial loads. We address these issues by using a centered-log transform on relative abundance data and including log-scaled bacterial load as a potential feature to select. While some bootstrapped models selected bacterial load as a positive predictor (Figure 5c), the majority of models did not. This further suggests that the majority of microbiome information is encoded in the relative ratios of taxa abundance, which is broadly consistent with previous findings [27,28].

Finally, our study is limited to a cross-sectional analysis, limiting us to making predictions on lung function state at the same time-point as microbiome sample and patient medical record collection. Assessing and refining our predictive machine learning algorithms on subsequent lung function data is an important future goal. Our primary objective is to predict future disease states and preemptively identify patients in need of medical intervention using early warning microbiome markers. To this effect, we plan to continue our analysis on a cohort of patients across time to evaluate predictive capacity for future health status.

We note that the major predictors identified in our models have been identified in various studies, and taken piecemeal there is less insight. The value of this work lies in the systematic integration of these multiple data sources (from both EMR and microbiome data sources). Our model comparisons (with / without EMR predictors) allow an assessment of the impact of oral bacteria, with and without key potential confounds. Ignoring these confounds could lead to spurious retention of microbiome taxa that correlate strongly with e.g. age or BMI. In addition, our analyses allows assessment of disparate factors on a common predictive scale – indicating for example that the impact of 1 standard deviation shift in *Fusobacterium* abundance is comparable to a 1 S.D. shift in BMI. Our model comparison approach lends more confidence to the conclusion that the retained oral taxa are associated with patient outcomes via causal pathways that are largely independent of age or BMI, being robust to their presence or absence in the predictive models. The research agenda of pursuing the nature of the causal pathways linking oral bacteria in the lung with patient outcomes is now on a firmer footing as a result of our study.

In summary, our study finds that inclusion of non-pathogenic taxa significantly improves model prediction accuracy of patient health status. We identify two oral-derived taxa (*Fusobacterium, Rothia*) that are independently informative of lung function, which may be either biomarkers or potential probiotics. Our results call attention to the potential predictive utility of oral microbes (regardless of their functional roles) in the clinical assessment of CF patient health.

## Methods

### Subjects

All procedures performed in studies involving human participants were in accordance with the ethical standards of the institutional and national research committees. Authorization was obtained from each patient enrolled according to the protocol approved by the Emory University Institutional Review Board (IRB00042577).

### Sample collection and 16S analysis

Expectorated sputum samples were obtained from CF patients attending the Children’s Healthcare of Atlanta and Emory University CF Care Center from January 2015 to August 2016. De-identified patient information including age, sex, height, BMI, CFTR genotype, degree of glucose control (HbA1c), and ppFEV1) were obtained (Table 1). Among these CF patients, 39 were diagnosed with CF-related diabetes patients (CFRD) by a CF endocrinologist. HbA1c value was missing for one CFRD subject.

All patients were clinically stable, defined as having no increase in respiratory symptoms compared to baseline, and no acute illness or new medication for three weeks prior to sputum collection. Upon collection, sputum samples were diluted 1:3 (mass:volume) with PBS supplemented with 50 mM EDTA. Diluted samples were then homogenized by being repeatedly drawn through a syringe and 18-gauge needle. The resulting sputum homogenates were aliquot and stored at −80 °C until all 77 samples were collected. Microbiology culture results were obtained for sputum samples sent to the Clinical Microbiology laboratory on the same day as samples for sequencing were collected.

DNA was purified from sputum homogenate with the MoBio Power Soil kit (MoBio, Carlsbad, CA). The 16S V4 region was amplified and sequenced using Illumina MiSeq, yielding an average of 137,708 sequences per sample. Sequences were quality filtered and amplicon sequence variants were obtained using the QIIME2 deblur plugin. Taxonomic assignments were classified against both SILVA and Greengenes 16S reference databases and assigned based on highest taxonomic resolution. To mitigate compositional effects, 16S data were center-log transformed prior to all analyses. Nucleotides are uploaded to BioProject accession no. PRJNA666192.

### Statistical and Quantitative Analysis

Patient samples were binned by ppFEV1 (Normal: >80%, Mild: 80-60%, Moderate: 60-40%, Severe: <40%). Variance across lung function categories in patient metadata and 16S metadata was tested using ANOVA.

Variation between microbiome composition and ppFEV1 was tested using Mantel tests on Bray-Curtis distances at 9999 permutations. Within-sample and among-sample diversity was calculated using the Shannon diversity index and Bray-Curtis based PCoA on 16S quantitation agglomerated to the genus level [40]. Associations between continuous variables were tested using Spearman correlations. A full pairwise correlation matrix was calculated, with rows and columns ordered by hierarchical clustering [41].

### Machine Learning

We use ElasticNet to fit regularized linear models predicting lung function (ppFEV1) from patient metadata, microbiome composition, and clinical microbiology results [42]. ElasticNet solves a penalized linear regression model using a weighted average of L1 (LASSO) and L2 (ridge regression) penalties. This limits overfitting by penalizing non-zero coefficients. We split our samples using a simple 70:30 train-test holdout, where models are trained on 53 samples and used to predict on the remaining 24. All input features were standardized (mean=0, S.D.=1) prior to model training to allow between-feature interpretability. From our full dataset, we create 4 additional data subsets: CF Pathogens, All 16S Data, Metadata, and Metadata + Pathogens. We include within-feature shuffling on the full set as a non-informative negative control.

We employ two methods to assess model robustness, and compare model performance using mean squared error (MSE). We generate 1000 bootstrap resampled sets from the training set and fit an ensemble of regularized linear models to obtain distributions for each model coefficient. We identify key metadata and taxa robustly selected (nonzero coefficients) across the ensemble of models. We assess model generalizability using leave-one-out cross-validation on the training set and compare resulting MSE ranges.

## Acknowledgements

We thank Karan Kapuria and Eunbi Park for help with bio-informatic pipeline development, and Peng Qiu for machine learning guidance. We acknowledge the technical support and provision of the clinical data by the CF Biospecimen Registry at the Children’s Healthcare of Atlanta and Emory University CF Discovery Core. This research was supported in part through research cyber-infrastructure resources and services provided by the Partnership for an Advanced Computing Environment (PACE) at Georgia Tech.

## Disclaimer

The funders had no role in study design, data collection and analysis, decision to publish, or preparation of the manuscript.

