## Supplemental for "Microbiome data enhances predictive models of lung function in people with cystic fibrosis"

**Supplementary Material**

*Detailed Sequencing Analysis*

Samples were sent to MR DNA Lab (Shallowater, TX) for DNA extraction, sequencing library preparation, Miseq sequencing, and absolute 16S quantitation. Microbiology culture results were obtained for sputum samples sent to the Clinical Microbiology laboratory on the same day as samples for sequencing were collected.

The V4 region of the resulting DNA was amplified with the 16S universal primers 515F (5'-GTGCCAGCMGCCGCGGTAA-3') and 806R (5'-GGACTACHVGGGTWTCTAAT-3'). A single-step 30 cycle PCR integrating sequencing amplification and library adapter/barcode attachment was performed using the HotStarTaq Plus Master Mix Kit (Qiagen, USA) by first incubation at 94 °C for 3 minutes, followed by 28 cycles of 94 °C for 30 seconds, 53 °C for 40 seconds and 72 °C for 1 minute, followed by a final elongation step at 72 °C for 5 minutes. Amplification products were then normalized, pooled and purified using calibrated Ampure XP beads for Illumina Miseq sequencing.

Illumina Miseq sequencing generated in a total of 10,603,544 sequences, with an average of 137,708 sequences per sample (minimum 76,281, maximum 191,868). All sequence processing was done through QIIME2 2018.2.0. Raw sequences were firstly de-multiplexed and quality filtered on a per-nucleotide basis (min quality: 4, window: 3, min length fraction: 0.75, max ambiguous: 0). Reads were denoised using the deblur plugin, and the sequences were trimmed at the length of 250 bp (sample stats: T, mean error: 0.005, indel_prob: 0.01, indel_max: 3, min_reads: 10, min_size: 2, jobs_to_start: 1). Taxonomic assignments were classified against both the SILVA and greengenes database and assigned based on their highest taxonomic resolution. Discrepancies were resolved manually through BLAST and comparing against the non-redundant NCBI sequence database.

Based on taxonomic information, microbiome composition data was obtained for every sputum sample and a phylogenetic tree was constructed via *fasttree*. To correct for the variation 16S rDNA copy number among different taxa, the number of sequences per sample were divided by known 16S rDNA copy number of the genus or divided by four (average number of 16S rDNA copy number) if the information was missing.[37] Samples were rarefied to 17000 reads to guarantee equal sampling for subsequent analysis.[38]

*Machine Learning*

To illustrate our machine learning approach, we begin with the model output trained on the full dataset (all 16S and metadata predictors, Figure S2). Figure S2a plots predicted versus observed lung function, for both the training dataset (data on 53 patients used to train model parameters) and the test dataset (data on 24 patients held back during model training). Figure S2b highlights the parameters retained in the predictive model and their weighting. Our initial machine learning analysis (Figure S2) suggest that the addition of non-pathogen 16S data improves model performance as evidenced by the retention of non-pathogen predictors in a penalized regression, and flags specific taxa as potential predictors.

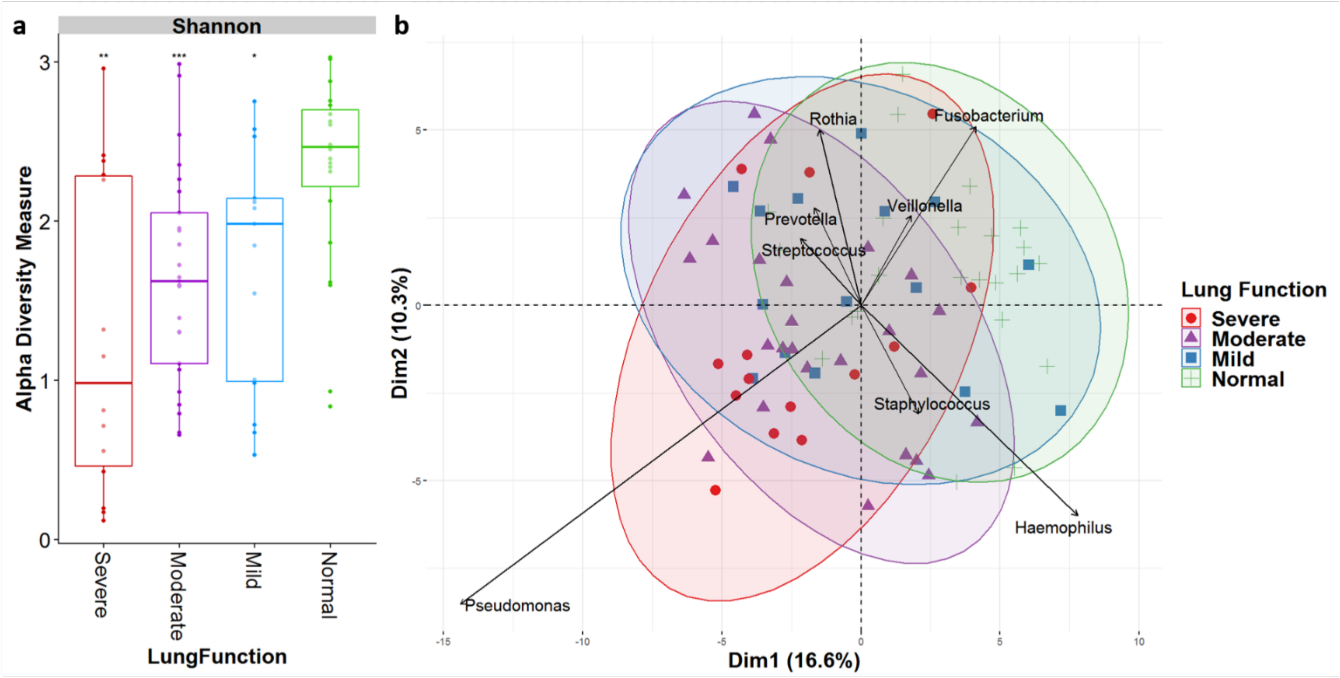


***Figure S1. Shannon diversity and ordination. a)*** *Within-sample diversity (Shannon index) is lower in severe disease states compared to normal (Kruskal-Wallis, p<0.01).* ***b)*** *Between-sample diversity (Bray-Curtis PCoA on top 25 genera, centered log-ratio transformed). PCs 1 and 2 combined explain ~27% of the microbiome variance, and weakly clusters patients by Lung Function.*

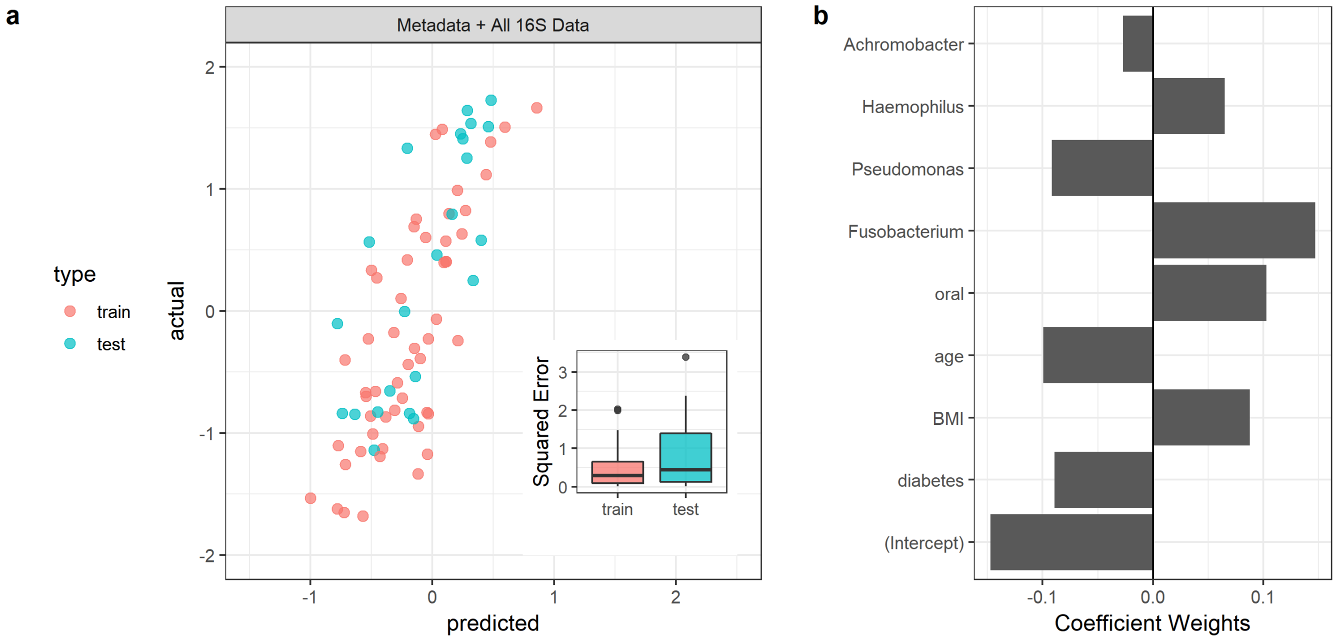


***Figure S2. ElasticNet-identified predictors of lung function.*** *We train a baseline predictive model of ppFEV1 using the ElasticNet algorithm (alpha = 0.5) to perform feature selection. We assess the train-test holdout method on metadata + all 16S data. The train-test uses a standard 70-30 split (53 patient training set, 24 patient test set). (**a**) We plot model-predicted ppFEV1 values (scaled) against actual ppFEV1 values and calculate squared errors for each data point. We find that the model trained with the full dataset has the highest performance (see Figure S1 for prediction subset model performance) and selects features across different input data sources. (**b**) Model coefficients from the train-test holdout show general agreement with CF heuristics. Age, diabetes, Achromobacter and Pseudomonas abundance are selected as negative predictors of age whereas Haemophilus, Fusobacterium, oral taxa abundance, and as BMI are positive predictors.*

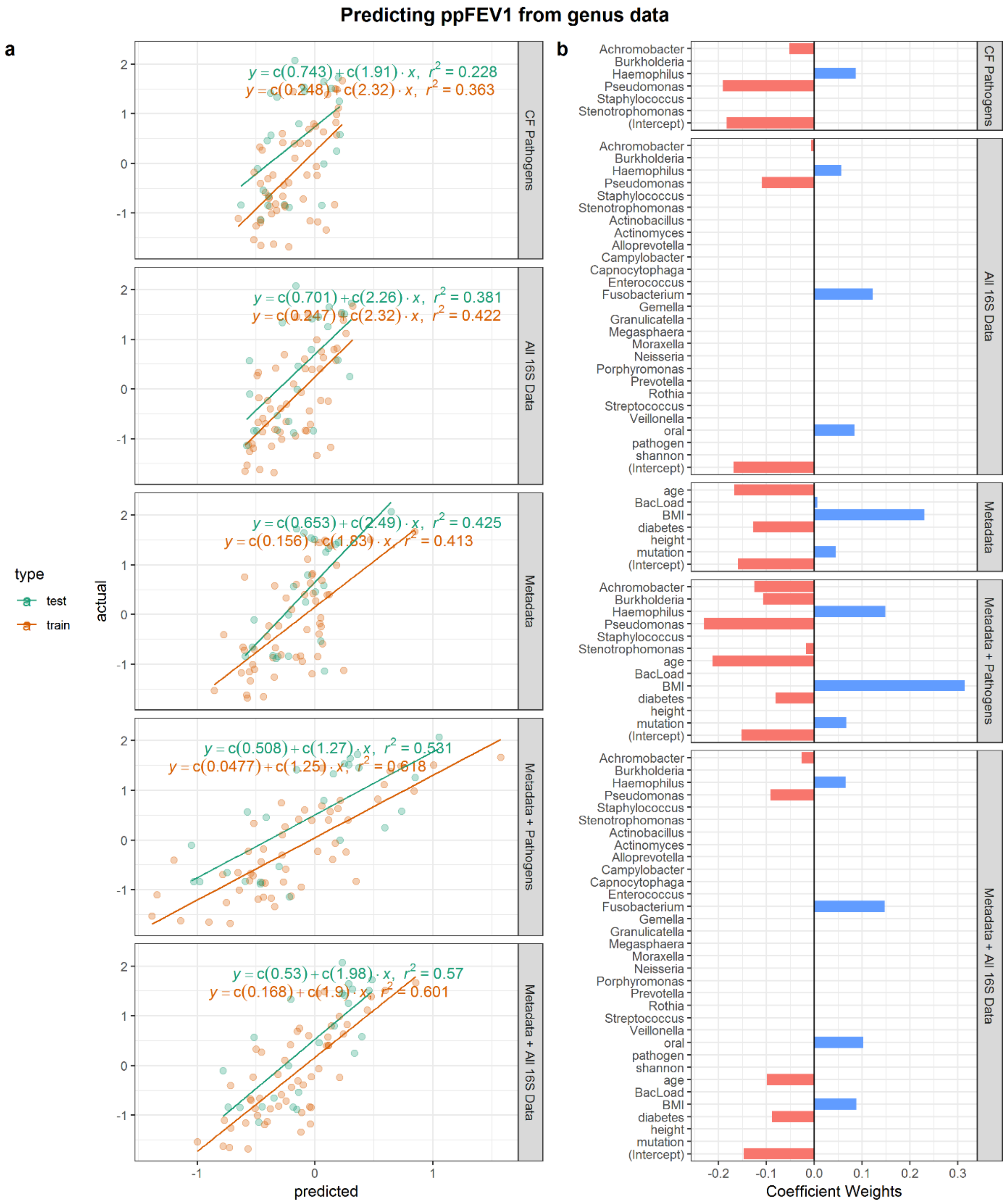


**Figure S3. Predicting ppFEV1 from genus data.** *To obtain baseline models, we assess the train-test holdout method on five input data sources: 16S quantitation of CF Pathogens (clr-transformed), all 16S data (clr-transformed), metadata, metadata + pathogens, and metadata + all 16S data.*

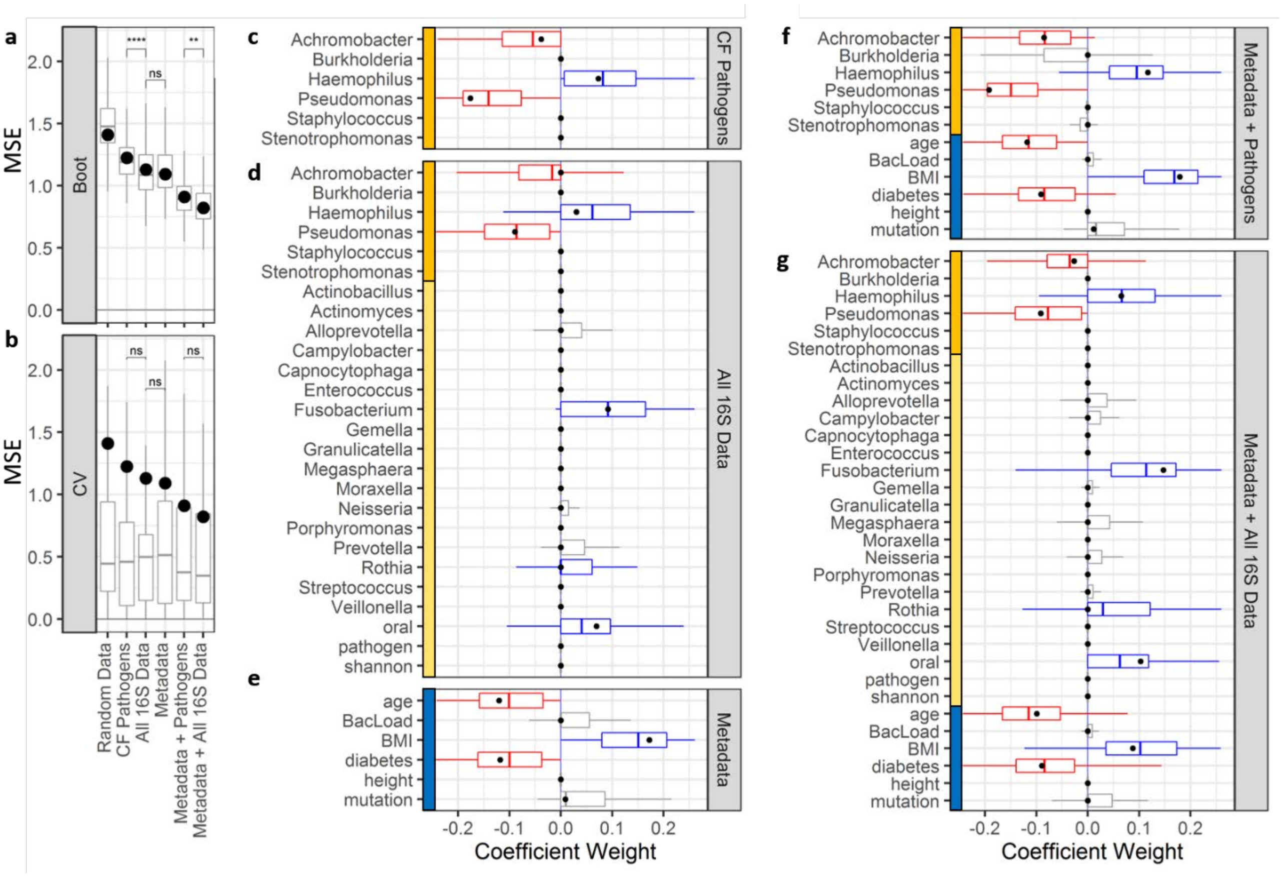


***Fig S4. Bootstrapped ElasticNet-identified predictors of lung function.*** *ML models were trained using varying input datasets.* ***a)*** *1000-fold bootstrapping and* ***b)*** *leave one out cross-validation (LOOCV) were used to generate prediction error (MSE) ranges across feature subsets. Models trained on all of the data show lower error compared to other feature subsets. Adding 16S pathogen quantitation decreases model error. Models trained on all 16S data outperform models using only 16S quantitation (p < 0.01, t test). Regardless of input features, models trained on the full sample set (black points) are greater than median LOOCV MSEs (boxplots).* ***c-g)*** *Coefficient ranges for train/test (black points) and bootstrapped models (boxplots) trained on varying input datasets (blue: metadata, orange: 16S pathogens, yellow: 16S other taxa) show consistency between both machine learning strategies. Both cases select* Pseudomonas *and* Achromobacter *as negative predictors*.
